# SNPector: SNP inspection tool for diagnosing gene pathogenicity and drug response in a naked sequence

**DOI:** 10.1101/834580

**Authors:** Peter T. Habib, Alsamman M. Alsamman, Ghada A. Shereif, Aladdin Hamwieh

## Abstract

Due to the ability to diagnose diseases early and evaluate the effectiveness of medicinal drugs, single nucleotide polymorphism (SNP) identification receives significant interest. Detection and diagnosis of genetic variation through skill-less computational tools would help researchers reducing the severity of such health complications and improving the well-tailored therapies using discovered and previously known information. We introduce SNPector, which is a standalone SNP inspection software could be used to diagnose gene pathogenicity and drug reaction in naked genomic sequences. It identifies and extracts gene-related SNPs, and reports their genomic position, associated phenotype disorder, associated diseases, linkage disequilibrium, in addition to various drug reaction information. SNPector detects and verifies the existence of an SNP in a given DNA sequence based on different clinically relevant SNP databases such as NCBI Clinvar database, Awesome database, and PharmGKB and generates highly informative visualizations of the recovered information.

## Introduction

In recent years, the number of cases of genetically originated diseases has increased, alarming the world and sparking interest in the development of precision medicine using molecular biomarkers. Single nucleotide polymorphism (SNP), the most common genetic difference among individuals, occurs in the human genome. These randomized modifications in DNA bases cause alterations in protein sequence residues of amino acids, thus altering their functions which lead to different disease conditions in individuals (1). Several of these SNPs have been identified as disease-related genetic markers that have been used to recognize genes responsible for a particular disease in humans (2)□.

Distinguishing the evidence and the interpretation of a rich range of markers will be necessary to relate the major alterations in the SNPs and to discover their connection in the progression of disease. Clarification of the phenotypic-associative mechanisms for these variations is therefore vital for comprehending the sub-atomic subtleties of disease start and for developing novel therapeutic methods (3),(4).

Although SNPs may exist in various areas of the gene, such as promoters, introns, 5′-and 3′ UTRs, to date, most research has focused on disease-associated SNPs (daSNPs) in coding regions or exons, especially non-synonymous SNPs, which may alter the biochemical ability of encoded proteins. In turn, altering gene promoters impact gene expression by changing transcription, binding transcription factor, methylation of DNA and modifications of histones. As a consequence, changes in gene expression, their impact on disease susceptibility, and drug responses can differ depending on the location of the SNP (5)–(7).

With the expansion of genetic variants, different software could be used to generate new knowledge to support disease diagnosis and drug response studies and to develop new biomarkers for disease identification and drug customization. In this regard, a number of software applications have been developed in the last few years to classify, prioritize and evaluate the impact of genomic variants.

For example, the Ensemble Variant Effect Predictor offers access to a large range of genomic annotations, with a variety of frameworks that answer different needs, with easy setup and evaluation methods (8)□. Similarly, SnpEff categorizes the results of genome sequence variations, annotate variants according to their genomic location and estimates the coding effects. Depending on genome annotation, it is possible to predict coding effects such as non-synonymous or synonymous substitution of amino acids, stop codon gains or losses,start codon gains or losses, or frame changes (9)□.

On the other hand, PolyPhen-2 assesses the potential impact of the genetic substitution of amino acids on the basis of physical, evolutionary comparative factors and model structural changes. Based on these profiles, the probability of a missene mutation becoming dangerous is measured on the basis of a combination of all these properties (10)□. In like manner, SIFT calculates whether the substitution of amino acids affects protein activity, based on the homology of sequences and the physical properties of amino acids. It may be used for non-synonymous polymorphisms and laboratory-induced missense mutations that naturally occur, to effectively classify the effects of SNPs as well as other types, including multiple nucleotide polymorphisms (MNPs) (11)□.

Moreover, Phyre2 is a web-based suite of tools for predicting and analyzing protein structure, function and mutations. It has sophisticated remote homology identification methods to build 3D models, anticipate ligand binding sites, and evaluate the effect of amino acid variants, e.g. non-synonymous SNPs (12)□. Missense 3D uses the user-provided UniProt ID of the query protein, wild-type residue and substitution and other information to generate PDB residue mapping and predict the substitution effect on the 3D protein structure (13)□.

To conclude the effect and possible phenotype of SNP, these software and web applications require minimum information such as SNP genomic position, SNP ID, allele form, and/or gene name. Acquiring these information require using different computational tools, extensive time and some analysis skills. Most of the time, only gene sequences are available in which the SNPs are hidden without any additional information.

Int this regard, we introduce SNPector, which is a standalone SNP inspection software could be used to diagnose gene pathogenicity and drug reaction in naked genomic sequences. It identifies and extracts gene-related SNPs, and reports their genomic position, associated phenotype disorder, associated diseases, linkage disequilibrium, in addition to various drug reaction information. It detects and verifies the existence of an SNP in a given DNA sequence based on different clinically relevant SNP databases such as NCBI Clinvar database (14)□, Awesome database (15)□, and PharmGKB (16)□. Lastly, it connects identified SNPs, related diseases and drugs, and produces numerous visualization figures to explain these relationships with the support of different Python modules.

## Design and implementation

SNPector was written using Pythpn3 programming language as a standalone package and could be run on different operating systems platforms supported with Python 3.x compilers. To achieve user-friendly usage, the SNPector can be operated from a console through simple command line (Figure 1).

SNPector use different SNPs record information collected from NCBI (159,184 record), Awesome (1,080,551 record), and PharmGKB (17) (3,932 record). Ldlink is an online tool can be used to assess linkage imbalance (LD) throughout ancestral populations and is a popular method to exploring population-specific genetic framework and functionally navigation disease susceptibility areas (18). An Application Program Interface (API) has been programmed to download an LDhap file containing linkage disequilibrium statistics and potentially functional variants for a query variant resulted from sequence.

SNPector starts by running BLAST (19) software locally to find out the genomic location of a given DNA sequence on human genome. If successfully, it retrieves SNPs record located within query genomic range using NCBI ClinVar database. According to retrieved records from database, the detected SNPs in user-provided queries are marked as wild or mutated. Additionally, more information regarding detecting SNPs records will be retrieved from different implemented databases. These information will be used to generate different illustration figures.

If process is successfully finished, SNPector will generate four different files; (A) Text file contains the output BLAST result, where the genomic location of the user-defined sequences is predicted, (B) Tab delimited file contains SNPs retrieved NCBI database located in the same regions, (C) Two files regarding these specific SNPs information retrieved from Awesome and PharmGKB databases, (D) different figures depicting SNPs with a similar mutation effect to the detected SNPs located on other genomic regions, SNP linkage disequilibrium, the relationship between SNP, drug, and phenotype (Figure 2).

**Figure 1:**
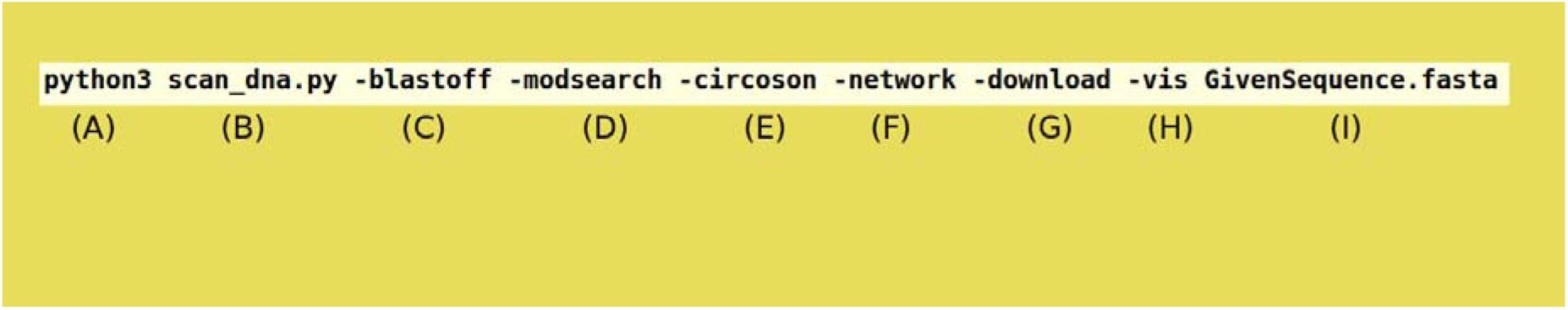
The SNPector command line structure. A) Python3 compiler, B) scan dna.py: The program main script, C) -blaston / -blastoff: in order to, initiate BLAST procedure for provided sequence against the genome to locate where the sequence is situated, if the blastoff is chosen the will use previous blast results, D) -modesearch / -modescan: Figure out SNPs that are located in the scope of query using different modes, E) -circoson : Draw circos figure to illustrate where SNP with same properties/effect are located, F) -networkon : in oreder to link between SNPs, diseases and drugs and produces network HTML file, G) -download: The activatation the API to download data for identified SNPs from LDlink database, H) -vis: in order to produce different figures and plots, I) GivenSequence.fasta : The user-provided sequence in fasta file format. Any of the previous parameter can be deactivated when replaced with -off.

## Results and discussion

SNPector can collect and retrieve information from the user-provided DNA sequence in the simplest way possible. By integrating different databases into SNPector, it is possible to detect the fluctuations in the abundance of SNPs in query through the comparison with known variants of human genome. Such steps are accompanied by the use of online and verified sources to gather previously published details regarding target genomic regions and to generate highly informative visualizations of the recovered information.

Many tools, however, provide SNPs annotation, but they are still limited to the information provided (Table 1). SNPector, on the other hand, provides a new technique that extracts SNP from a naked sequence with no prior information. In addition, other benefit of SNPector is to annotate the discovered SNPs based on various known databases. Moreover, SNPector provides user with more deeper and visualization figures, highlighting other SNPs with similar mutation effect on protein phosphorylation, ubiquitination, methylation, or sumoylation sites, and predicts substrates of N-acetyltransferase.

Additionally, SNPector provides the ability to visualize obtained information about the linkage disequilibrium of detected SNPs using various python packages such as Matplotlib (20), generating a number of figures that summarize vast amounts of previously published data indicating SNPs allelic segregation, association, minor allele frequency. Figure 2) shows an example of illustrations that can be generated through SNPector.

**Table 1:**
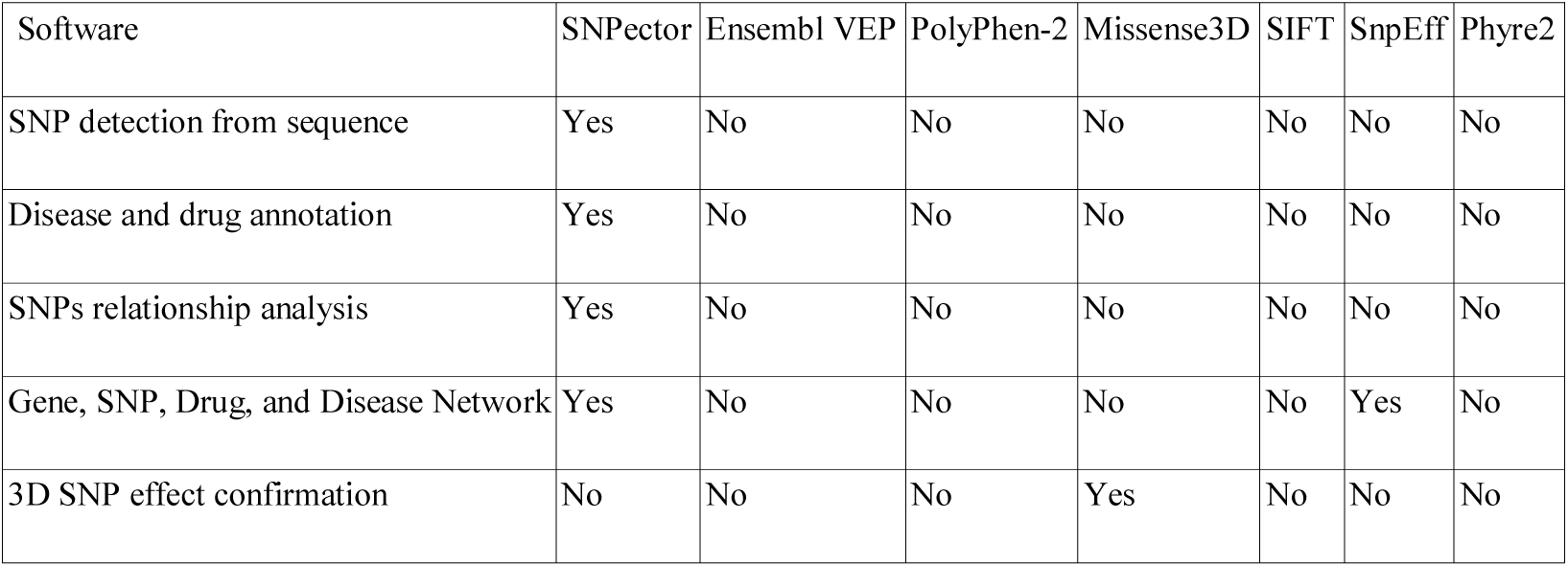
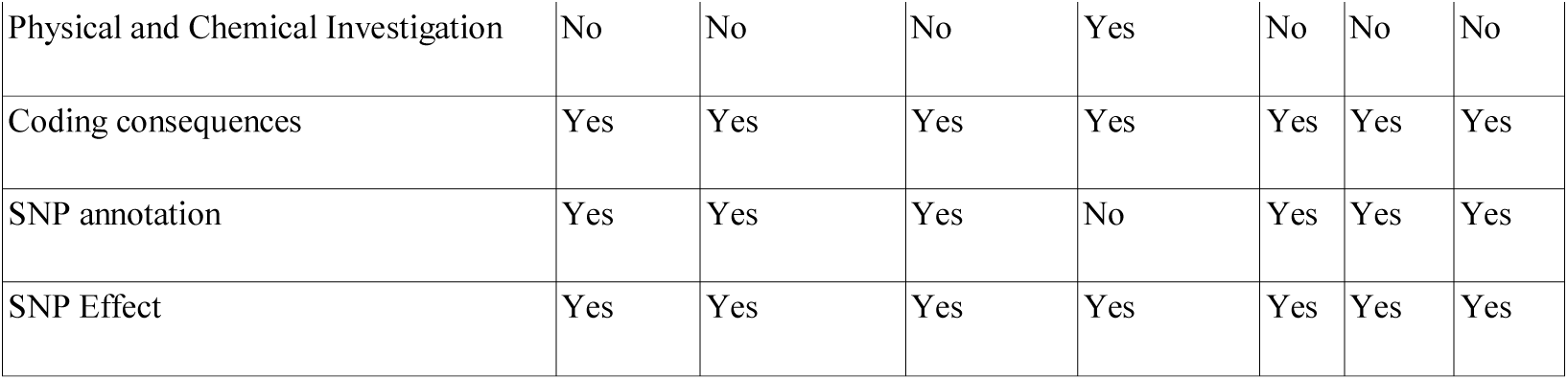
Comparison between SNPector and published SNP annotation tool.

**Figure 2:**
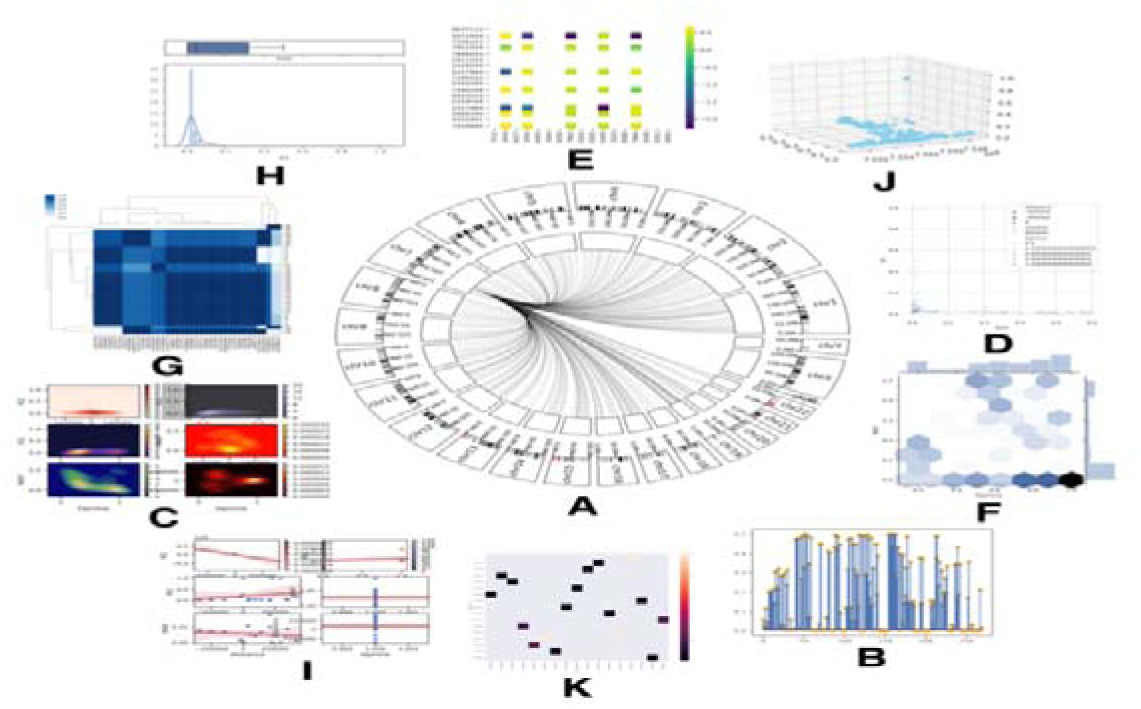
(A) Circos illustrate where other SNPs that have same proprieties are located. (B) Lollipop that show values each with head to be more distinguishable specially in lower values. (C) Counter Plot between two value creating this colored shade in which more contrast means higher value. (D) Numerical schematic show the distribution between four values by plotting and scaling color contrast according to other to values. (E) Heat map between SNP linkage disequilibrium matrix to show how each two SNPs are linked. (F) Marginal plot combine between column graph and plot both show the relationship between two values. (G) Dendogram with heatmap which show how far all SNP are linked to each others. (H) Histogram with box plot to compare visually between two values. (I) Plotting illustrate the regression fit of Two plotted value. (J) 3D plot of three values. (K) Annotated heatmap show the plotted value with its number on it with changing in color contrast according to the number.

## Conclusion

One of the currently growing medical research paradigms is the diagnosis of genetic virulence that accumulates in our genome causing catastrophic health problems. Detection and diagnosis of genetic variation through skill-less computational tools would help researchers reducing the severity of such health complications and improving the well-tailored therapies using discovered and previously known information.

SNPector provides and detects all available information about the disease-related SNPs in the given query with minimum user-provided information. It connects between different available information and produce various illustrations depicting SNP related diseases and treatment network, linked disequilibrium, minor allele frequency, similar SNPs with the same mutation effect and other information.

---

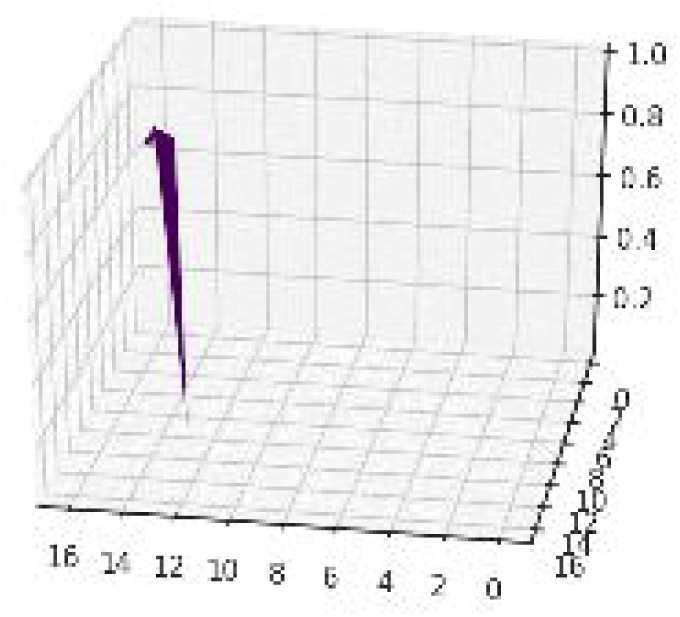

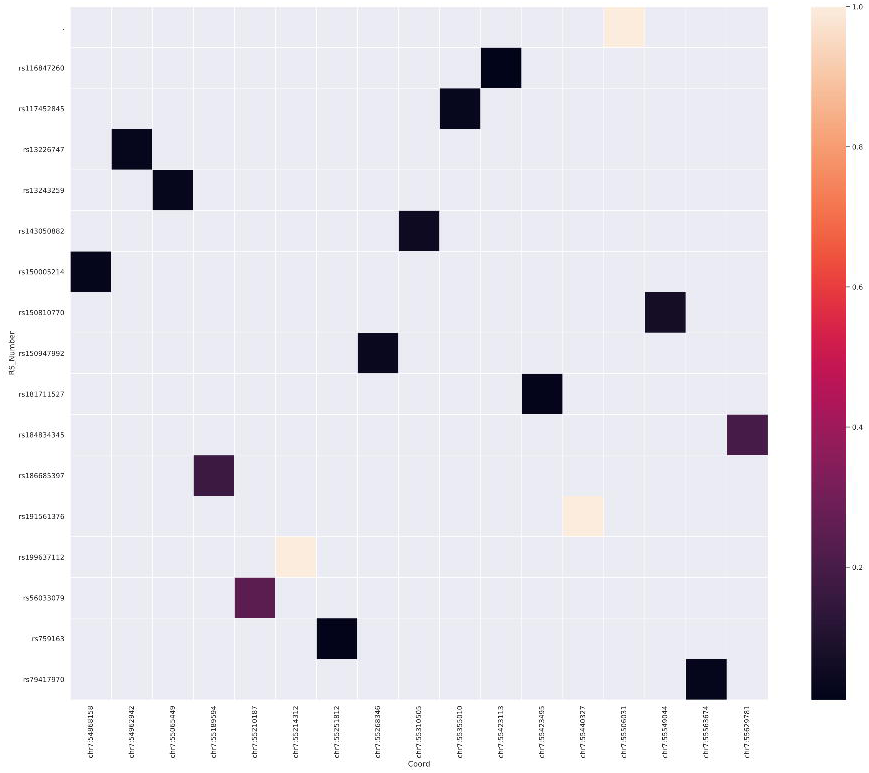

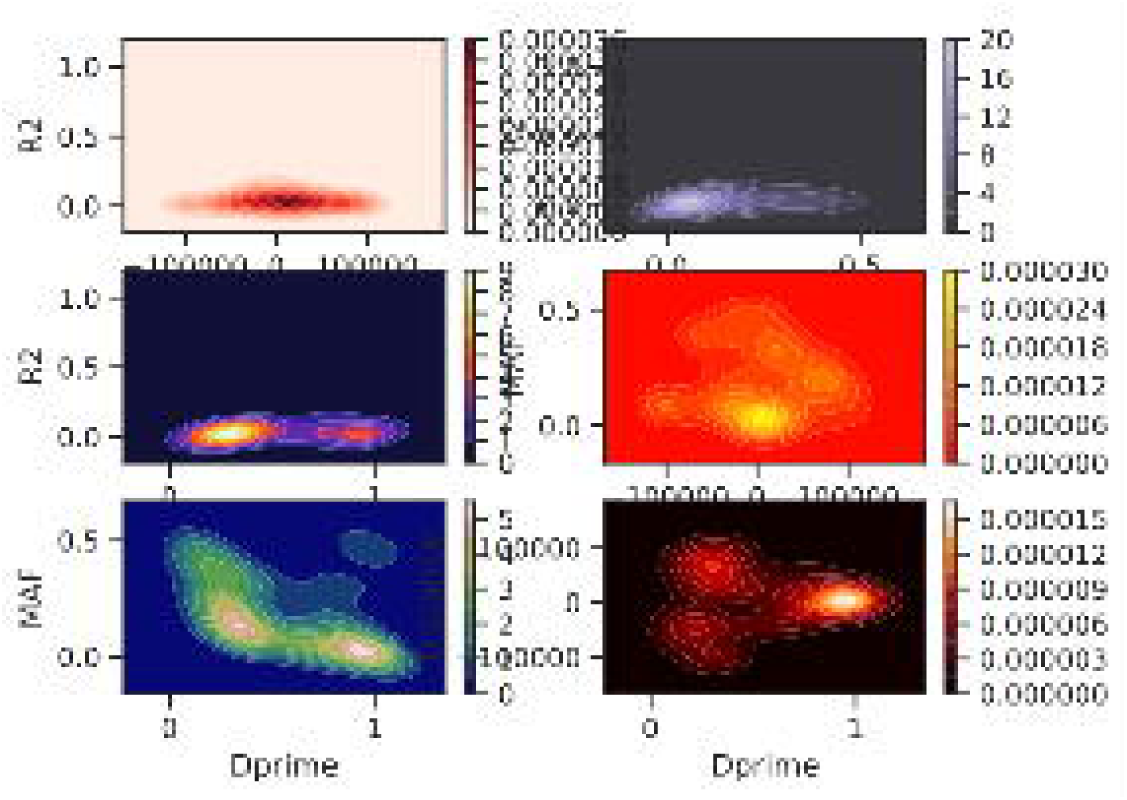

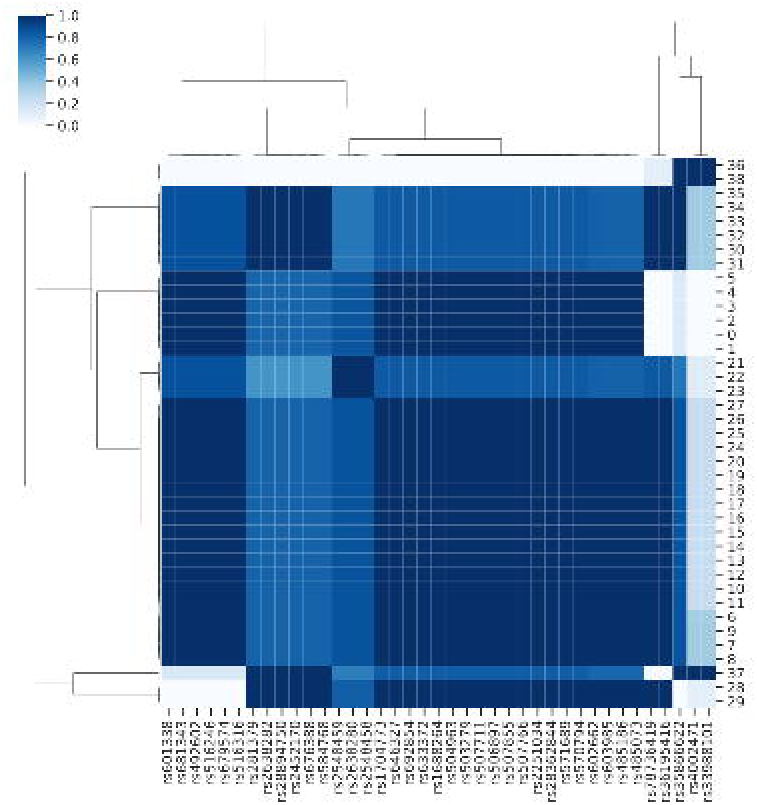

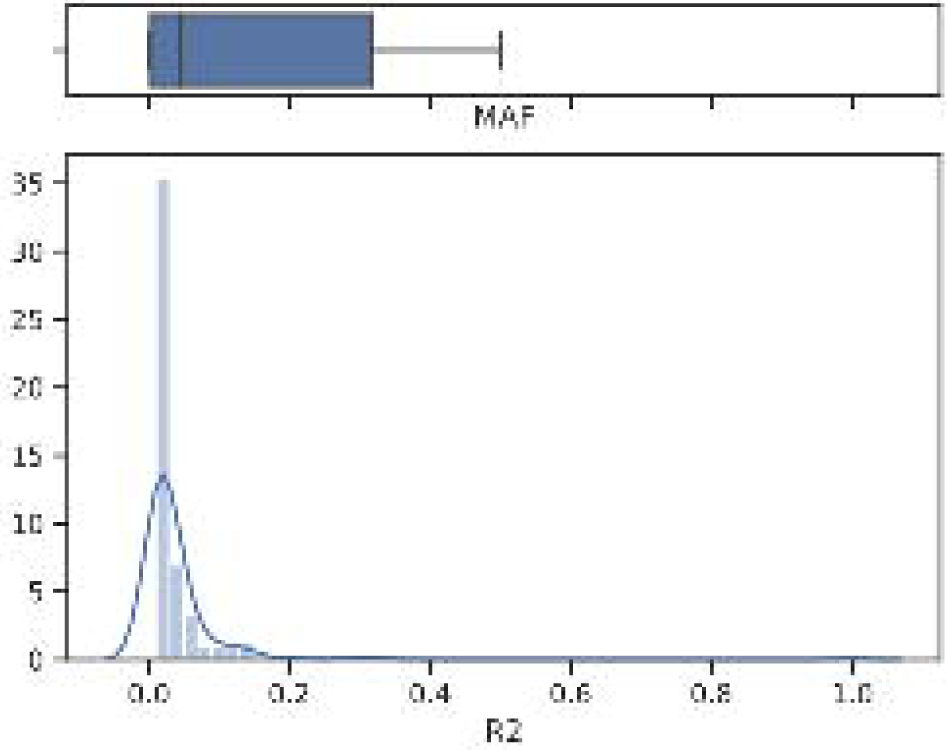

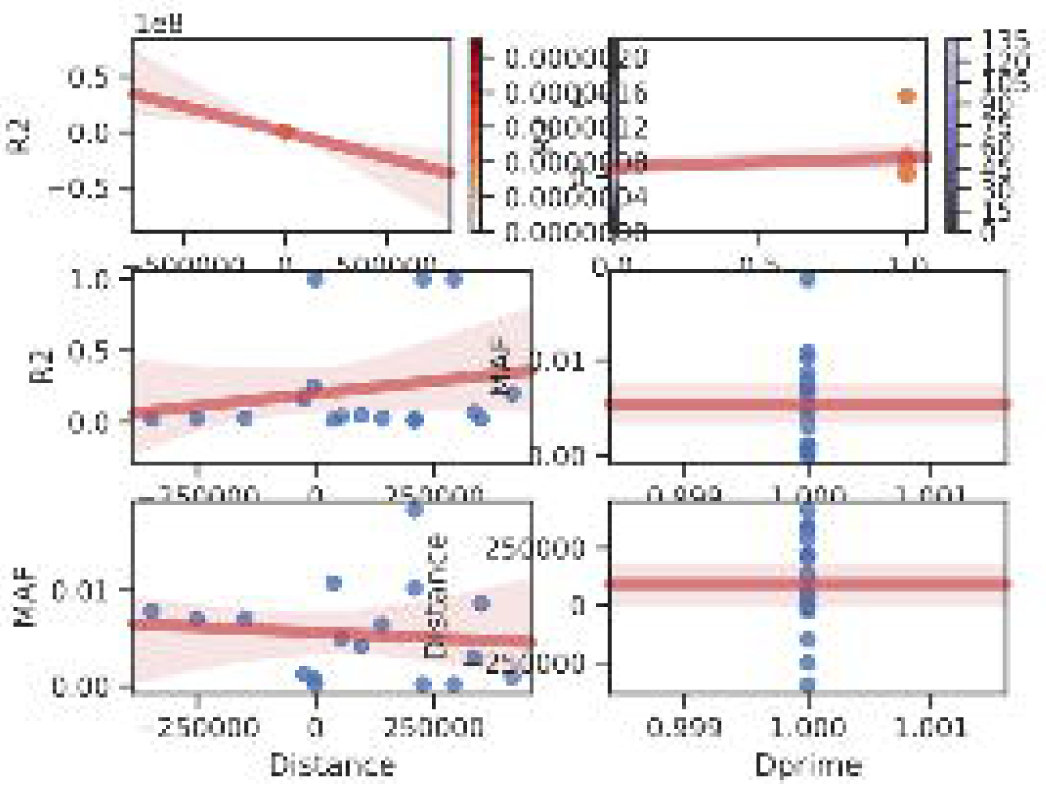

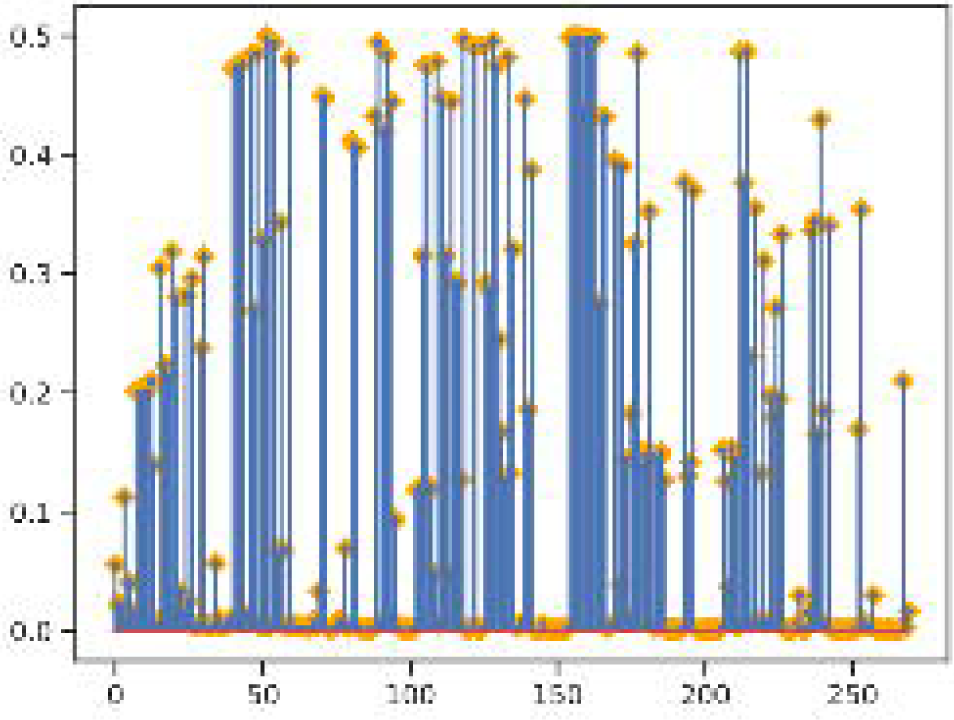

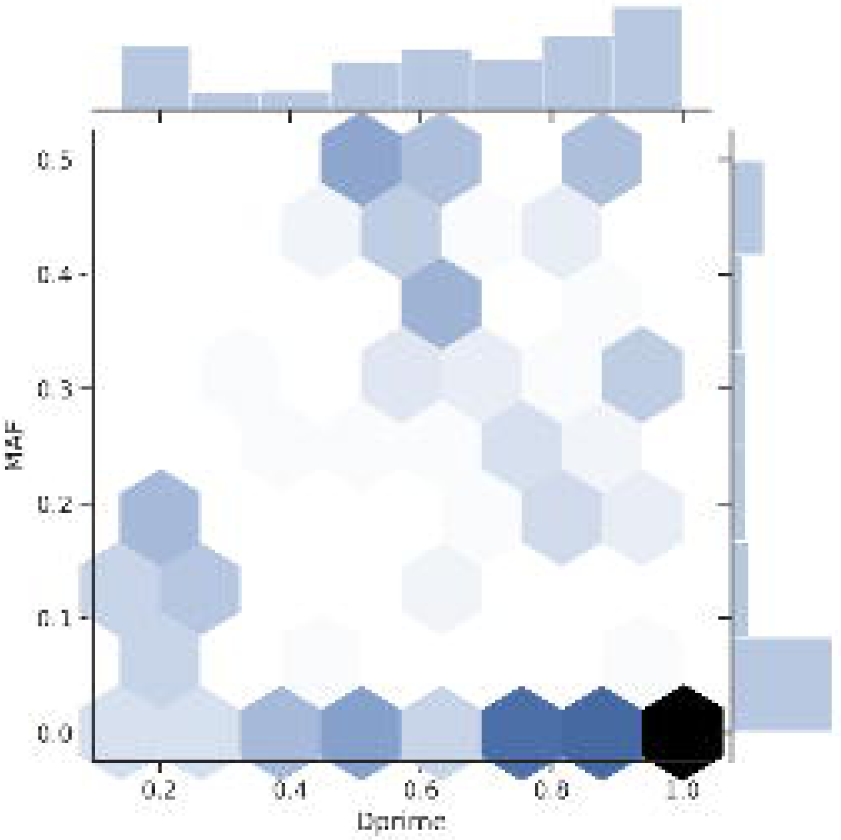

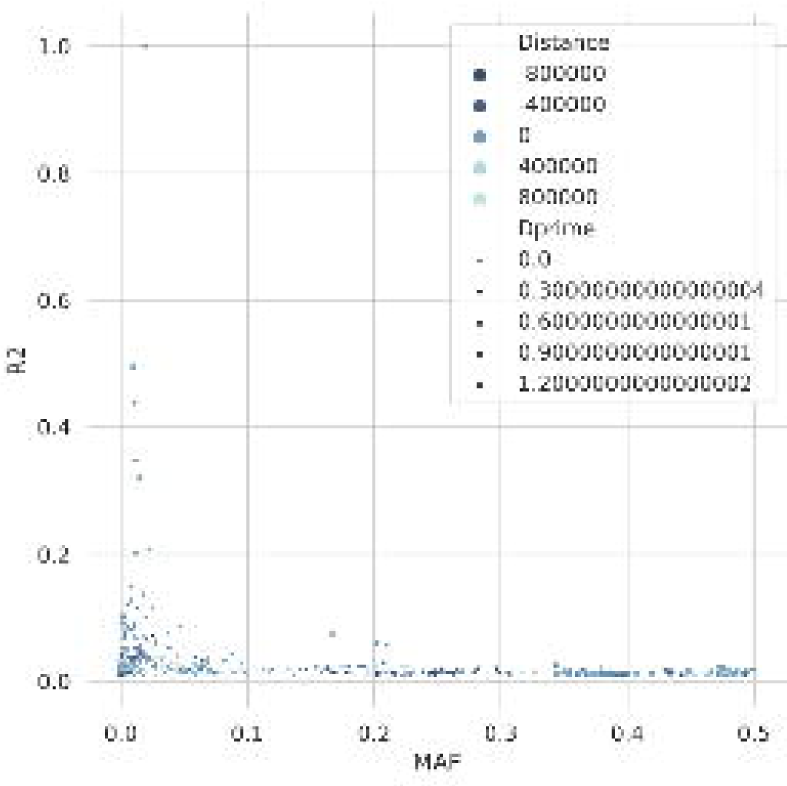

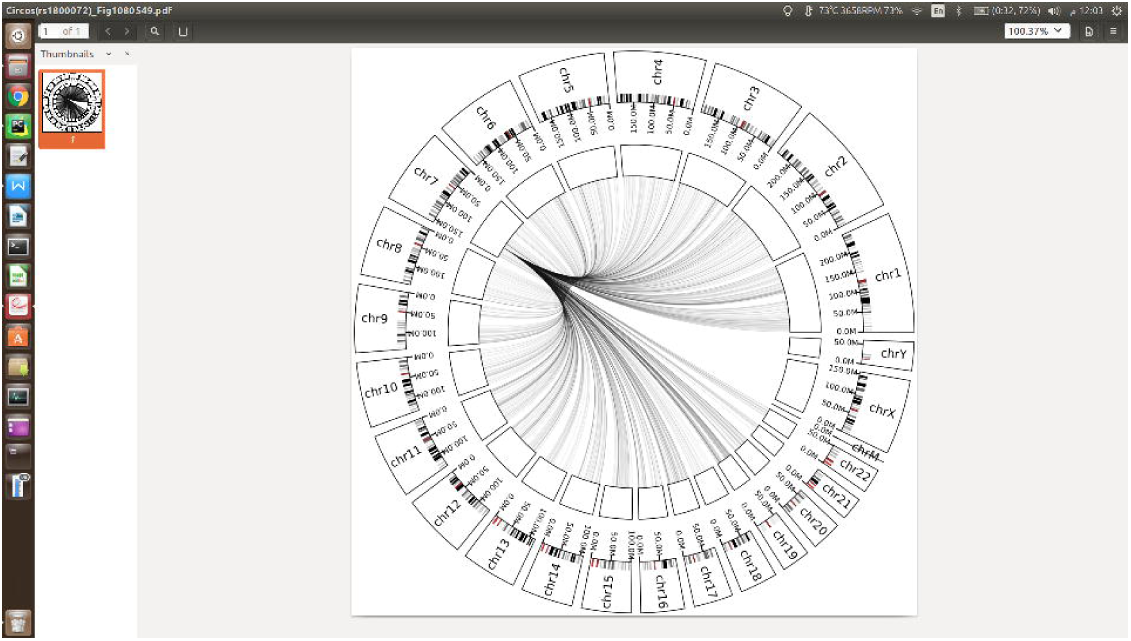

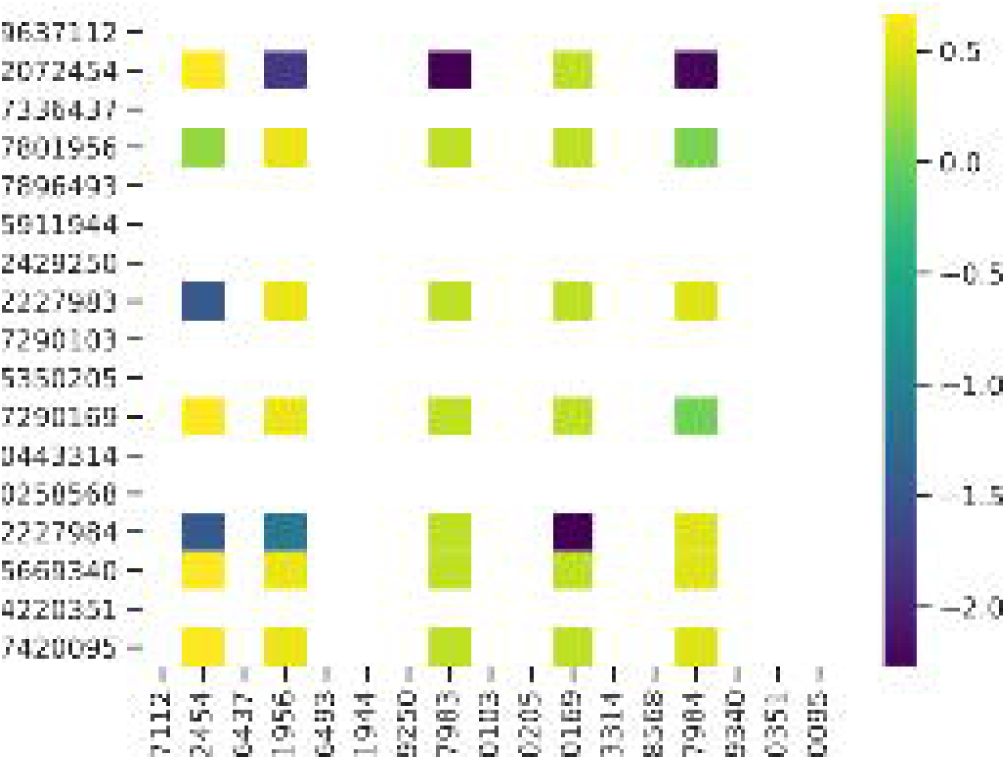

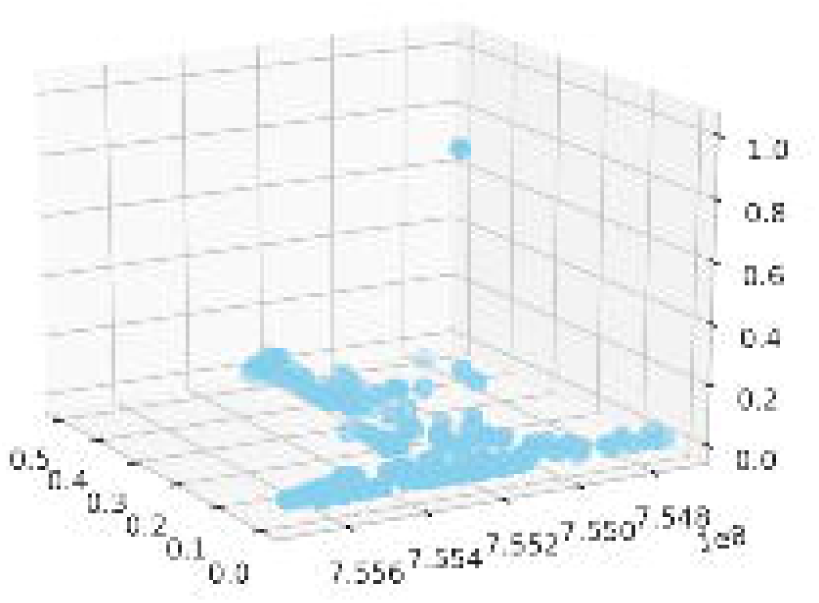

